# Visualization of peptidoglycan layer isolated from gliding diderm bacteria, *Flavobacterium johnsoniae* and *Myxococcus xanthus*, by quick-freeze deep-etch replica electron microscopy

**DOI:** 10.1101/2025.01.30.635643

**Authors:** Yuhei O tahara, Tâm Mignot, Makoto Miyata

## Abstract

The bacterial peptidoglycan layer plays an important role in protecting the bacteria from turgor pressure, viruses, and predators. However, it also acts as a barrier in transmitting forces generated on the cell membrane to adhesion proteins on the surface during gliding locomotion. In this study, peptidoglycan layers were isolated from two species of gliding diderm, i.e., Gram-negative bacteria, and their structures were visualized by quick-freeze deep-etch replica electron microscopy. The diameter of pores in the peptidoglycan layer of *M. xanthus* and the area of surface pores were 51 nm and 14.6%, respectively, which were significantly larger than those of *E. coli* (32 nm and 5.8%) and *F. johnsoniae* (29 nm and 7.0%). Based on this, we discussed the mechanism by which diderm bacteria transmit forces across the PG layer to the bacterial surface.

**Significance:** Peptidoglycan is a rigid meshwork that protects bacterial cells from turgor stress, virus, and predators. Bacteria move to get nutrients and escape from predators. The force for motility is generated in the cell membrane and transmitted to the surface across the peptidoglycan layer. In this study, the peptidoglycan structures of two representative gliding Gram-negative bacteria were visualized to understand the transmission mechanism.

## Introduction

Peptidoglycan (PG) is a common peripheral structure in bacteria that contributes to shape maintenance and turgor resistance of the cell body [1]. It is a filament composed of two amino sugar units, N-acetylglucosamine (GlcNAc) and N-acetylmuramic acid (MurNAc), crosslinked laterally by peptides to form a meshwork. As PG is a bag-like structure that covers the entire cell, its reorganization is necessary during cell growth and division [2]. The PG mesh can allow macromolecules such as proteins to pass through, however large complexes larger than 5 nm are blocked [3].

Many bacteria move to acquire nutrients or new genes or to escape predators or waste products and most bacterial motility is caused by flagellar rotation or type IV pili (T4P) retraction [4,5]. The proximal ends of flagella and pili are connected to force-generating units, which are embedded in the cell membrane. The force generated by the units in the membrane must be transmitted across the PG and, in the case of Gram-negative bacteria (which are also called “diderm”), further across the outer membrane to the cell surface. As these force-generating units partially penetrate the peptidoglycan and cell membrane, the forces generated inside the cell are directly transmitted to the outside [4,6,7]. However, the gliding motility of *Bacteroidota*, Gram-negative bacteria, and the T4P-independent gliding motility, “A motility” of *Myxococcus* require that the force generated in the cell membrane to be transmitted across the PG layer to a substrate such as glass or agar [8-10]. To discuss how forces are transmitted across the PG in these systems, information about the structure of the PG is essential. The information is: Are the PG layers of the board horizontally bonded? How hard is the PG layer? How hard is the PG layer? We have previously developed a method to isolate PG layers and reveal their detailed structures, and have visualized PGs of *Escherichia coli* and *Bacillus subtilis*, where PGs are isolated by treating cells with SDS and Proteinase K, and then observed by quick-freeze replica electron microscopy (EM) [11-14]. In the quick-freeze replica electron microscopy, samples are frozen within milliseconds using liquid helium. The sample is fractured with a knife, water evaporated, replicated with platinum, and then the replica is observed [15]. This method produces a high contrast image with a resolution that is much better than scanning electron microscopy (SEM). Other advantages of this method are low distortion due to mechanical force and high contrast, unlike AFM and cryo-electron microscopy [2,16]. In this study, we isolated and visualized PGs from non flagellated *E. coli, F. johnsoniae*, and *M. xanthus*, and analyzed and compared their structures in order to understand their motility mechanisms.

## Materials and Methods Strains and culture conditions

*E. coli* (DH5α) was cultivated in Luria-Bertani (LB) medium at 37°C with shaking at 180 rpm. *F. johnsoniae* and *M. xanthus* were grown in Casitone Yeast Extract (CYE) medium at 30°C with shaking at 150 rpm shaking [17,18]. Cells were harvested at an optical density of around 0.9 at 600 nm.

### Microscopy

Light microscopy was performed using a BX50 microscope with 100× objective and phase contrast optics (Evident, Tokyo, Japan) [11,19]. Electron microscopy was performed as previously described [11,12,19]. Image J ver1.52a was used for analysis.

### PG isolation

PG was isolated as previously described. Briefly, 1 mL of culture was collected by centrifugation at an OD_600_ around 0.9. Cells were suspended with 0.5 mL PBS (75 mM sodium phosphate (pH 7.3), 68 mM NaCl). Sodium dodecyl sulphate was added to be 0.5% and incubated at 96 °C for 3 h. PG was collected by centrifugation at 20,000 ×g for 20 min, and suspended in 1 mL PBS. This procedure was repeated twice. To the suspension, 10 μL of 10 mg/mL chymotrypsin was added and incubated at 37 °C for 2 h. The PG was collected by the centrifugation, suspended in 1 mL water and this process was repeated twice. The final suspension was made with 10 μL water.

## Results

### Visualization of peripheral structures including peptidoglycan layer

To obtain an overview of the peripheral structures, non flagellated *E. coli, F. johnsoniae*, and *M. xanthus* were observed by quick-freeze replica electron microscopy (Fig. 1). For *E. coli*, two types of images were observed: one in which the outer membrane and periplasmic layer were removed by fracturing, and the other in which the cytoplasm and other cross sections were exposed (Fig. 1A, D, E). We identified each layer, based on the thickness of each layer observed in the cross section and the order of the layers from the outside. The thickness of the periplasmic space was 28 ± 1.8 nm, thicker at the cell poles, in agreement with was consistent with previous reports. The surface of the PG was porous, which was observed in the area where the outer membrane and periplasmic space were removed. The cell membrane underlying the PG showed general undulation, large holes in some places, and small particles 7.5 ± 0.2 nm in diameter. The holes should be structures left after membrane proteins have been pulled out by the fracture, and the particles should be membrane proteins. The fractured images of *F. johnsoniae* showed that the outer membrane was detached in many cells, exposing the PG. The periplasmic thickness was 26.9 ± 0.7 nm, in agreement with previous reports (Fig. 1B). Fine pores similar to those of *E. coli* were observed on the PG surface below the outer membrane. The fractured images of *M. xanthus* showed that the outer membrane was detached, exposing the PG, as observed in *F. johnsoniae*. The periplasmic thickness was 28.2 ± 1.0 nm, in agreement with previous reports. No significant difference in periplasmic thickness was found between the three species (ANOVA p>0.5).

**Fig. 1.**
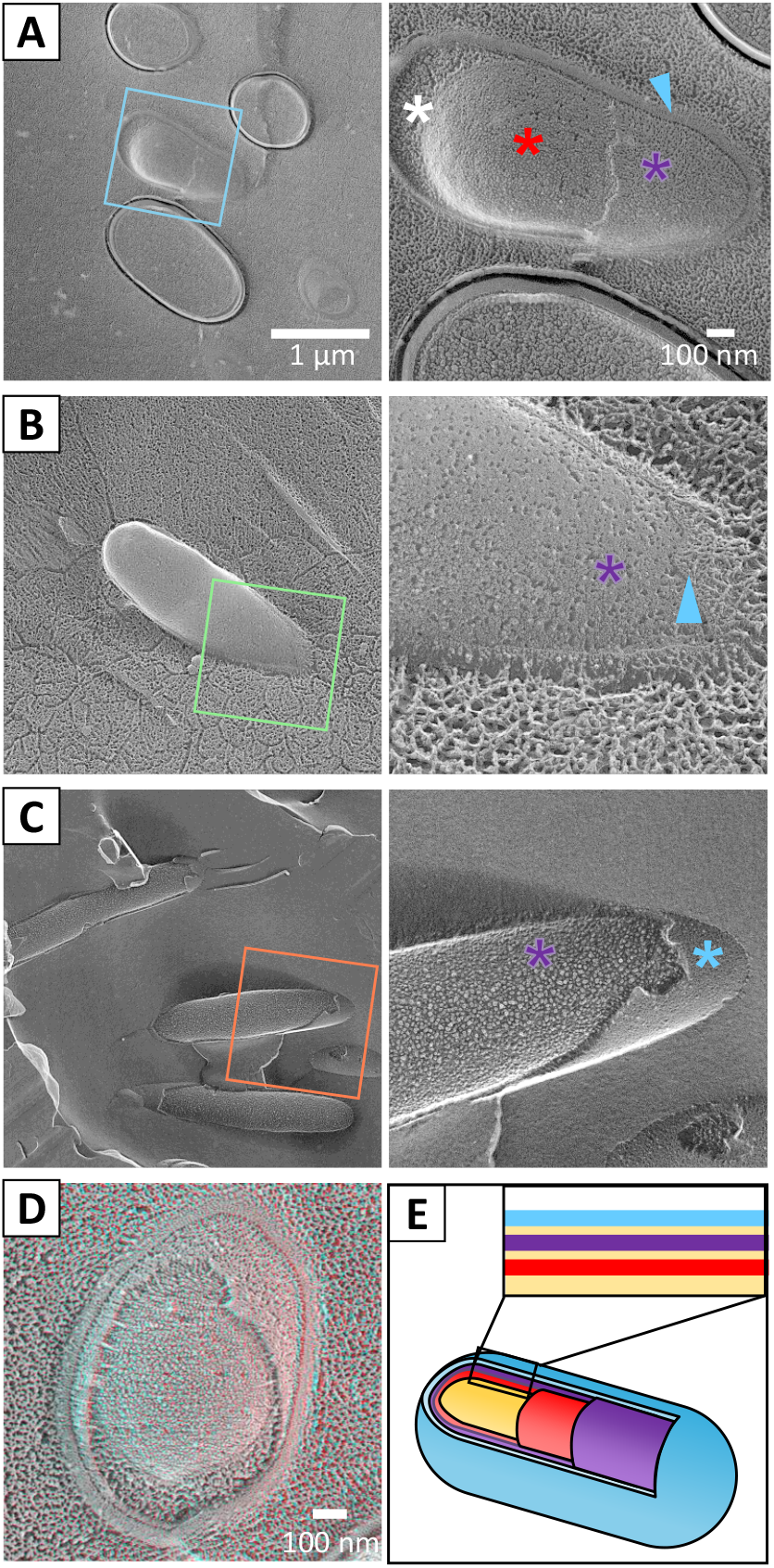
Freeze fracture image of bacterial cell membrane layers. Peripheral structures of three bacterial species visualized by freeze-fracture electron microscopy. Boxed areas in the left panel are magnified as in the right panel. Outer membrane, PG layer, and inner membrane are indicated marked by blue, purple and red colors and symbols, respectively. (A) *E. coli*. (B) *F. johnsoniae*. (C) *M. xanthus*. (D) Stereo view of a fractured *E. coli* cell. This can be observed through a pair of glasses with a red filter on the left side and a blue filter on the right side. For the stereo view, images of a field were taken at an angle of +7 degrees and −7 degrees, converted to red and blue, and superimposed [36]. (E) Cell schematic for the peripheral structure of Diderm (gram-negative) bacteria. Top: whole cell image. Bottom: section focused on peripheral structure.

### Isolation of peptidoglycan layers

We isolated PG layers from three bacterial species (Fig. 2). Light microscopy and negative staining EM images showed a clear loss of contrast in the isolated PGs in all three species, indicating that the treatment removed the membrane and cytoplasm. No significant damage was observed in the sac structures visualized by negative staining EM, indicating that sacs composed of PG layers could be isolated in an almost intact state. The results for *E. coli* were in agreement with previous reports. The widths of cells and isolated PG sacs measured by negative staining EM were 819.0 ± 16.8 nm and 1367.6 ± 46.0 nm, 506.1 ± 16.2 nm and 648.0 ± 21.3 nm, 426 ± 6.4 nm and 801.0 ± 58.0 nm for *E. coli, F. johnsoniae*, and *M. xanthus*, respectively (Fig. 2E). The width of the cell and PG sac could be explained by the relationship that the contents of the PG sac are removed from and it is flat, suggesting that the mesh-like PG sac is not significantly stretched when it covers the cell.

**Fig. 2.**
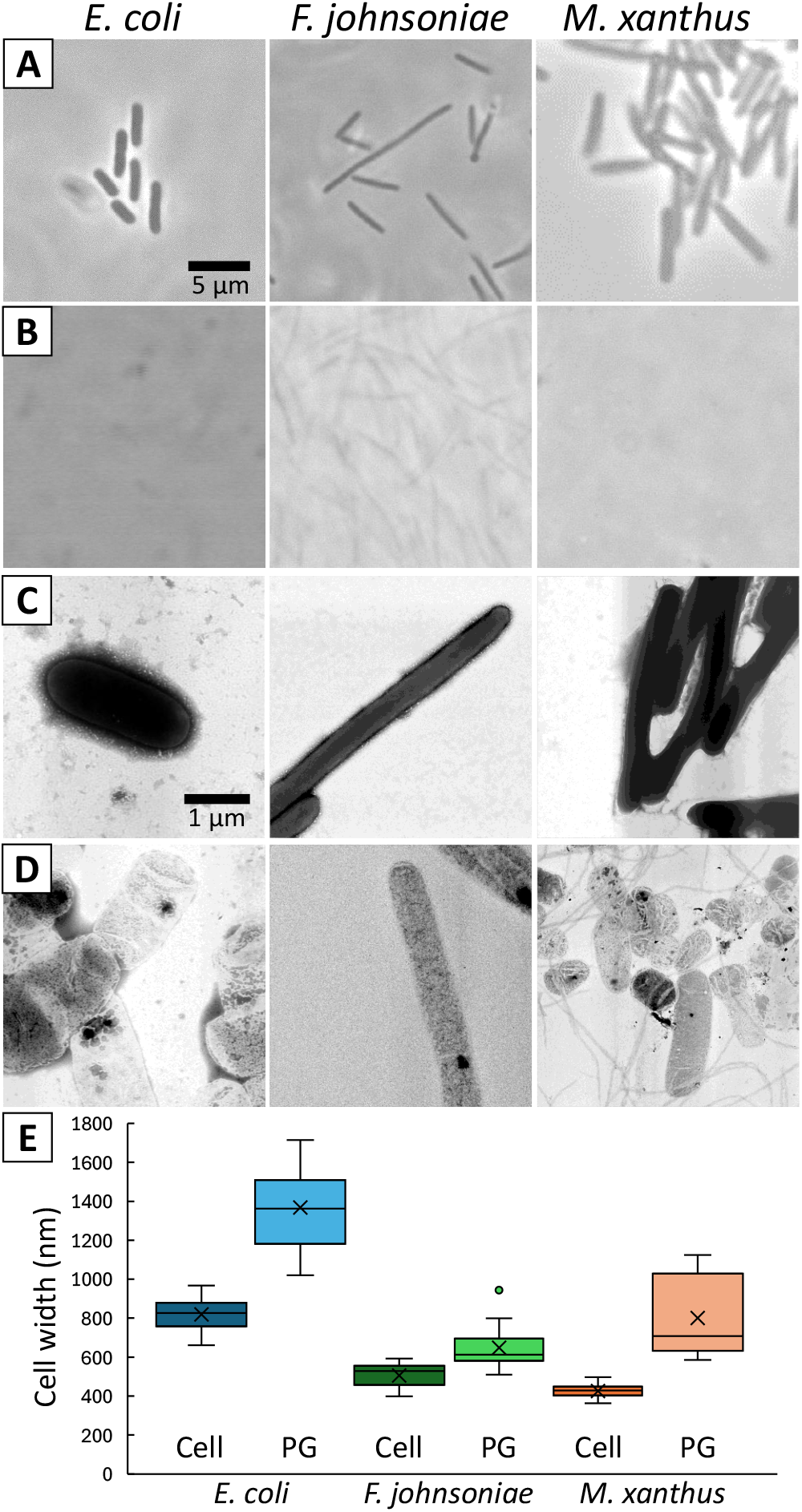
Optical and electron microscope images of purified peptidoglycan. PG layers isolated from three bacterial species. *E. coli, F. johnsoniae*, and *M. xanthus* are shown in the left, middle, and right columns, respectively. Phase contrast microscopy of live cells. (B) Phase contrast microscopy of isolated PG layers. (C) Negatively stained EM of cells. (D) Negatively stained EM of isolated PGs. (E) The widths of cells and isolated PG sacs measured by negative staining EM.

### Visualization of isolated PG by QFDE-EM

We observed and analyzed PG sacs isolated from each of the three bacterial species using QFDE-EM (Fig. 3). Most of the images obtained by this method also showed a sac shape, and no significant damage was found. The holes were more clearly visualized than by QFDE-EM of the cells or by negative staining of the isolated sac. The structure of *E. coli* was not significantly different from that of another strain previously published by our group. The holes were present throughout the PG layer sac, with no difference in size between the pole and the center. No directionality with respect to the cell axis was observed in the shape or orientation of the pores, which was also supported by the FFT (Fig. 3B).

**Fig. 3.**
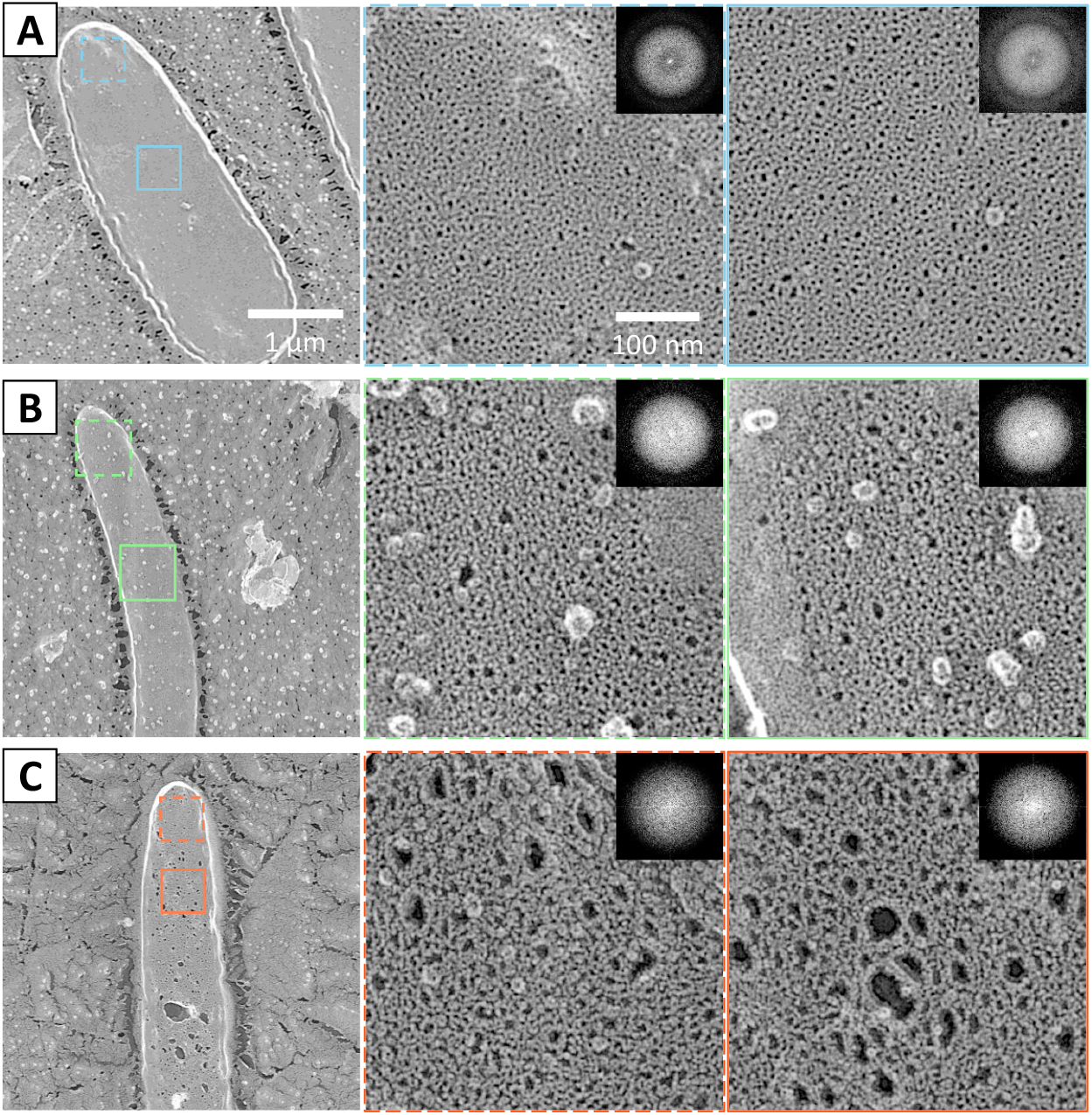
QFDE image of purified PG of *E. coli*. QFDE images of PG isolated from three bacterial species. The upper and lower boxed areas in the left panel are magnified as middle and right panels, respectively. The FFT image of the boxed area is shown in the upper right. (A) *E. coli*. (B) *F. johnsoniae*. (C) *M. xanthus*.

No significant damage was observed in the PG sac of *F. johnsoniae*, and the shape of the pore was clearly observed. No orientation of the pore shape or alignment with respect to the cell axis was observed. The PG layer of *M. xanthus* also retained the sac shape, and the pore shape was clearly observed as in the other two species. However, in some places the pores were significantly larger than others, indicating partial damage. Overall, the pores appeared larger than those of the other two species.

### Quantification of pore sizes in isolated PG sac

We quantified pore size to objectively characterize the three PG layers. We cut out areas of uniform distribution of holes from the PG sac images, binarized them, and judged the areas of high contrast between 10 nm^2^ and 1000 nm^2^ as ‘holes’ (Fig. 4A). We selected five cells from each species and summed these numbers (Fig. 6B). The number of holes detected in an area is shown in a violin plot for each species. The mean area of the holes was 31.8 ± 0.3 nm^2^ for *E. coli* (N= 8666); 29.1 ± 0.3 nm^2^ (N=8500) for *F. johnsoniae*, 51.2 ± 0.8 nm^2^ (N=8912) for *M. xanthus*. No significant difference in pore size was found between *E. coli* and *F. johnsoniae* (test: 0.226< p). However, the pore size of *M. thansus* was significantly different from that of *E. coli* and *F. johnsoniae* (test: p<0.0006), i.e. larger sized holes were more common in *M. xanthus* than in the other two species. We compared the percentage of area occupied by the holes in the area (Fig. 4C). The mean values were 5.8% ± 0.6% for *E. coli*, 7.0% ± 0.7% for *F. johnsoniae*, and 14.6% ± 1% for *M. xanthus* (N=5).

**Fig. 4.**
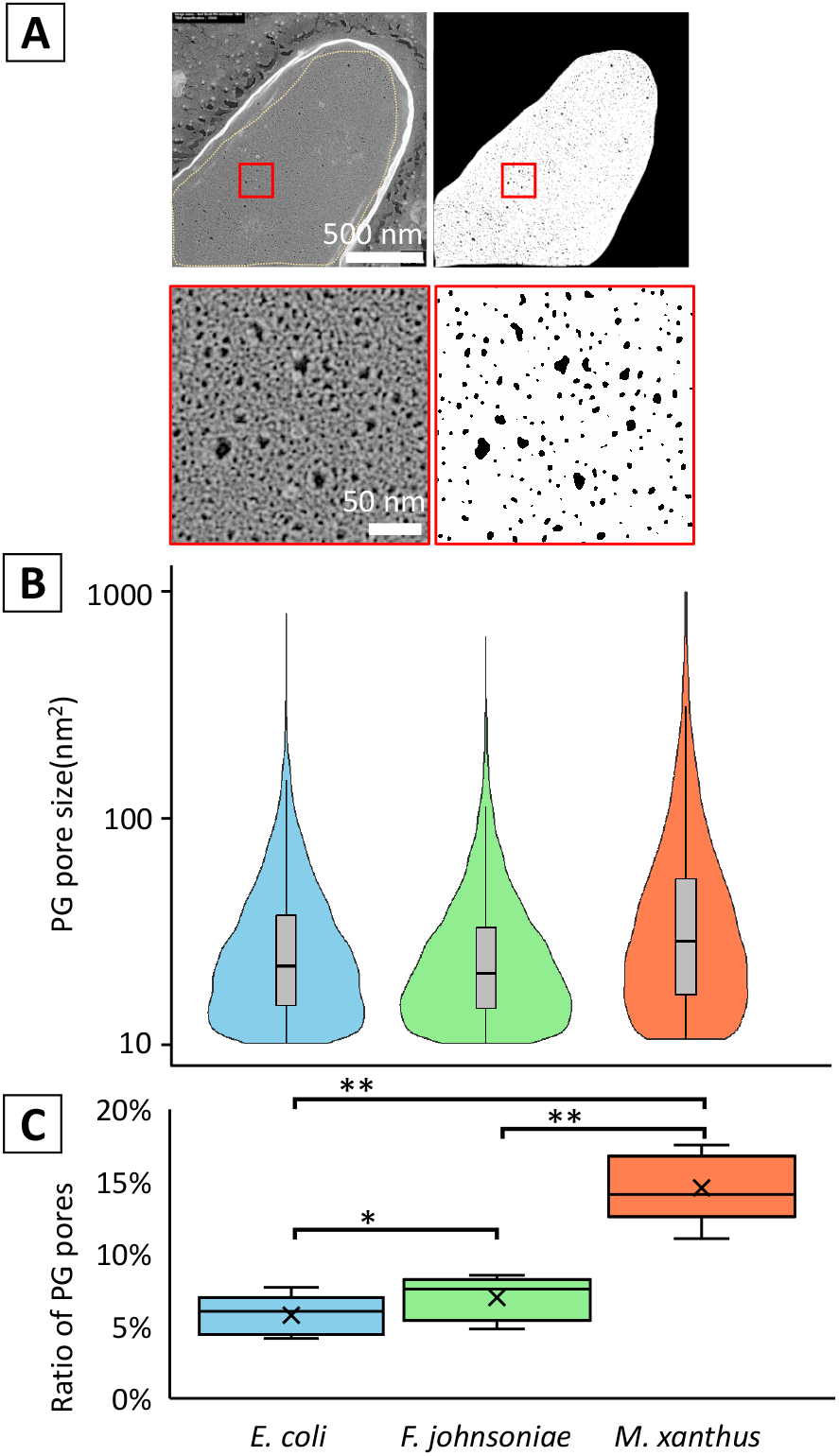
Analysis of pores in replica images of purified PG. Size distribution of PG pores. (A) Pore detection method. Images in the left panels were binarized as shown in the right panels. The boxed areas in the top panels are magnified in the bottom panels. (B) Violin plot and box plot overlaid with the size and number of pores. (C) Box plot of the percentage of pores in the area of PG. (ANOVA: P<1.3e-05, t-test: *P>0.2 ,**P<0.0005)

No significant difference between *E. coli* and *F. johnsoniae* (test: 0.226< p), while *M. xanthus* was significantly different from the other two species(test: p<0.0006).The Fourier transform of the image did not show any spots or distortions indicating periodicity in the shape of the holes.

## Discussion

The network structure of the *F. johnsoniae* PG sac was not significantly different from that of *E. coli* (Figs. 3 and 4), which is consistent with the fact that *F. johnsoniae* cells appear rigid under the light microscope, as do *E. coli* cells.

Force generation in *F. johnsoniae* is thought to be caused by proton flow through the inner membrane complex of GldL and GldM [8,20,21]. This complex is structurally similar to the stator of the flagellar motor. *F. johnsoniae* is observed to rotate in a precise circle around a position to which it adheres and clings to the glass surface, as observed as “tethered” in the bacterial flagellar motor [21-24], In addition, the adhesion protein SprB is anchored to the conveyor belt structure under the OM, and the conveyor belt structure is thought to move in a direction approximately along the cell axis by the proton gradient [25,26]. Based on these observations, a scenario is inferred in which the rotational motion generated by static GldL motors drive the conveyor belt (Fig. 5) [8,20,25]. If we accept this scenario, then there is no inconsistency with the PG structure of *F. johnsoniae* being common to *E. coli*, since the only motion transmitted from the cell membrane across the PG layer to the OM surface is the rotation of GldM within a motor complex like a shaft.

**Fig. 5.**
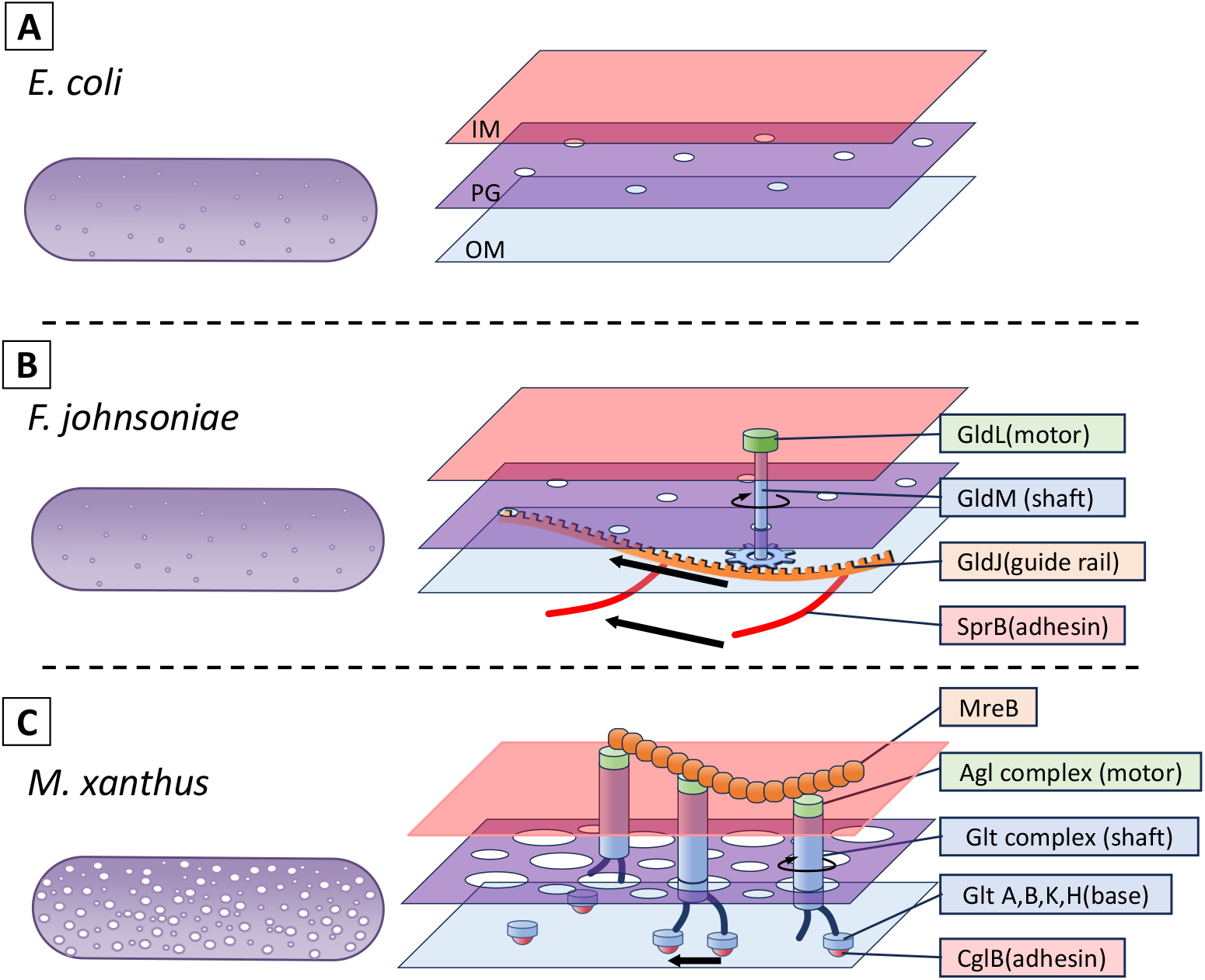
Model of PG in each bacteria. Schematic representation of PG and gliding machinery in Gram-negative gliding bacteria. On the left and right, the whole PG image and the enlarged PG image with gliding machinery are shown. The inner membrane (IM, cell membrane), the peptidoglycan (PG) layer, and the outer membrane (OM) are colored orange, purple, and right blue, respectively. (A) Non flagellated *E. coli. F. johnsoniae*. The static GldL generates mechanical torque through GldM periplasmic domain, which transmits across PG and the outer membrane like a shaft. The shaft rotation drives the belt composed of GldJ bound by the adhesion protein, SprB. (C) The Agl protein complex moves due to the proton motive force along an unknown intracellular track, possibly MreB. The motions are transmitted to an adhesion complex composed of the OM proteins Glt A, B,K, H and the surface exposed adhesin, CglB.

In *M. xanthus*, the area of each pore of the PG sac was larger than in *E. coli*, as was the percentage of the total pore area on the surface (Figs. 3 and 5). This may be one reason why *M. xanthus* cells are softer and have a much lower bending modulus than other bacteria [27]. We found a PG sac of *M. xanthus* with damage that was rarely observed in the other two species. This may also indicate that the PG sac of *M. xanthus* is softer and more fragile due to its porosity. *M. xanthus* has two different gliding motility mechanisms (Fig. 5) [28]. One is pili motility, which is observed in many bacteria such as *Pseudomonas, Neisseria*, and *Synechococcus*, and is also called “social motility” in *M. xanthus*. The other is “adventurous motility” characteristic of *M. xanthus*. This motility is also thought to be driven by a protein closely related to the stator of the flagellar motor, AglR, which generates force through a proton motive force [29-31].

However, unlike flagella and *F. johnsoniae*, this protein remains membrane-embedded and moves along an unknown track in a nearly axial direction. The fact that the adhesion part moves with the motor means that the lateral movement is transmitted across the PG layer into the extracellular space [9,17,28,32]. If the PG is a rigid sheet [12,16,33,34], forces exerted by the motor must be transmitted across the PG to the outside. How this occurs remains to be discovered, but a recent structural model predicts that flexible periplasmic domains of the motor-associated periplasmic Glt proteins link the IM and OM, connecting dynamically to an OM complex containing an adhesin (CglB) [17,35]. The PG holes observed in this study could accommodate these interactions and explain the transfer mechanism.

## Author Contributions

YOT did experiments and analyses. YOT and MM wrote a draft, discussed the story, and completed the manuscript.

## Data Availability Statements

The data underlying this article is presented in this article.

## Acknowledgements

We thank to Daisuke Nakane at the University of Electro-Communications for helpful discussions and Junko Shiomi at Osaka Metropolitan University for technical assistance. This study was supported by Grants-in-aid for scientific research (A) (JP17H01544), a JST CREST grant (JPMJCR19S5) to MM.

